# Phosphoryl group wires stabilize pathological tau fibrils as revealed by multiple quantum spin counting NMR

**DOI:** 10.1101/2024.08.14.606685

**Authors:** Lokeswara Rao Potnuru, Austin DuBose, Mesopotamia S. Nowotarski, Michael Vigers, Boqin Zhang, Chung-Ta Han, Songi Han

## Abstract

Hyperphosphorylation of the protein tau is one of the biomarkers of neurodegenerative diseases in the category of tauopathies. However, the molecular level, mechanistic, role of this common post-translational modification (PTM) in enhancing or reducing the aggregation propensity of tau is unclear, especially considering that combinatorial phosphorylation of multiple sites can have complex, non-additive, effects on tau protein aggregation. Since tau proteins stack in register and parallel to elongate into pathological fibrils, phosphoryl groups from adjacent tau strands with 4.8 Å separation must find an energetically favorable spatial arrangement. At first glance, this appears to be an unfavorable configuration due to the proximity of negative charges between phosphate groups from adjacent neighboring tau fibrils. However, this study tests a counterhypothesis that phosphoryl groups within the fibril core-forming segments favorably assemble into highly ordered, hydrogen-bonded, one-dimensionally extended wires under biologically relevant conditions. We selected two phosphorylation sites associated with neurodegeneration, serine 305 (S305^p^) and tyrosine 310 (Y310^p^), on a model tau peptide jR2R3-P301L (tau295-313) spanning the R2/R3 splice junction of tau, that readily aggregate into a fibril with characteristics of a seed-competent mini prion. Using multiple quantum spin counting (MQ-SC) by ^31^P solid-state NMR of phosphorylated jR2R3-P301L tau peptide fibrils, enhanced by dynamic nuclear polarization, we find that at least six phosphorous spins must neatly arrange in 1D within fibrils or in 2D within a protofibril to yield the experimentally observed MQ-coherence orders of four. We found that S305^p^ stabilizes the tau fibrils and leads to more seeding-competent fibrils compared to jR2R3 P301L or Y310^p^. This study introduces a new concept that phosphorylation of residues within a core forming tau segment can mechanically facilitate fibril registry and stability due a hitherto unrecognized role of phosphoryl groups to form highly ordered, extended, 1D wires that stabilize pathological tau fibrils.

**Graphical Abstract:** 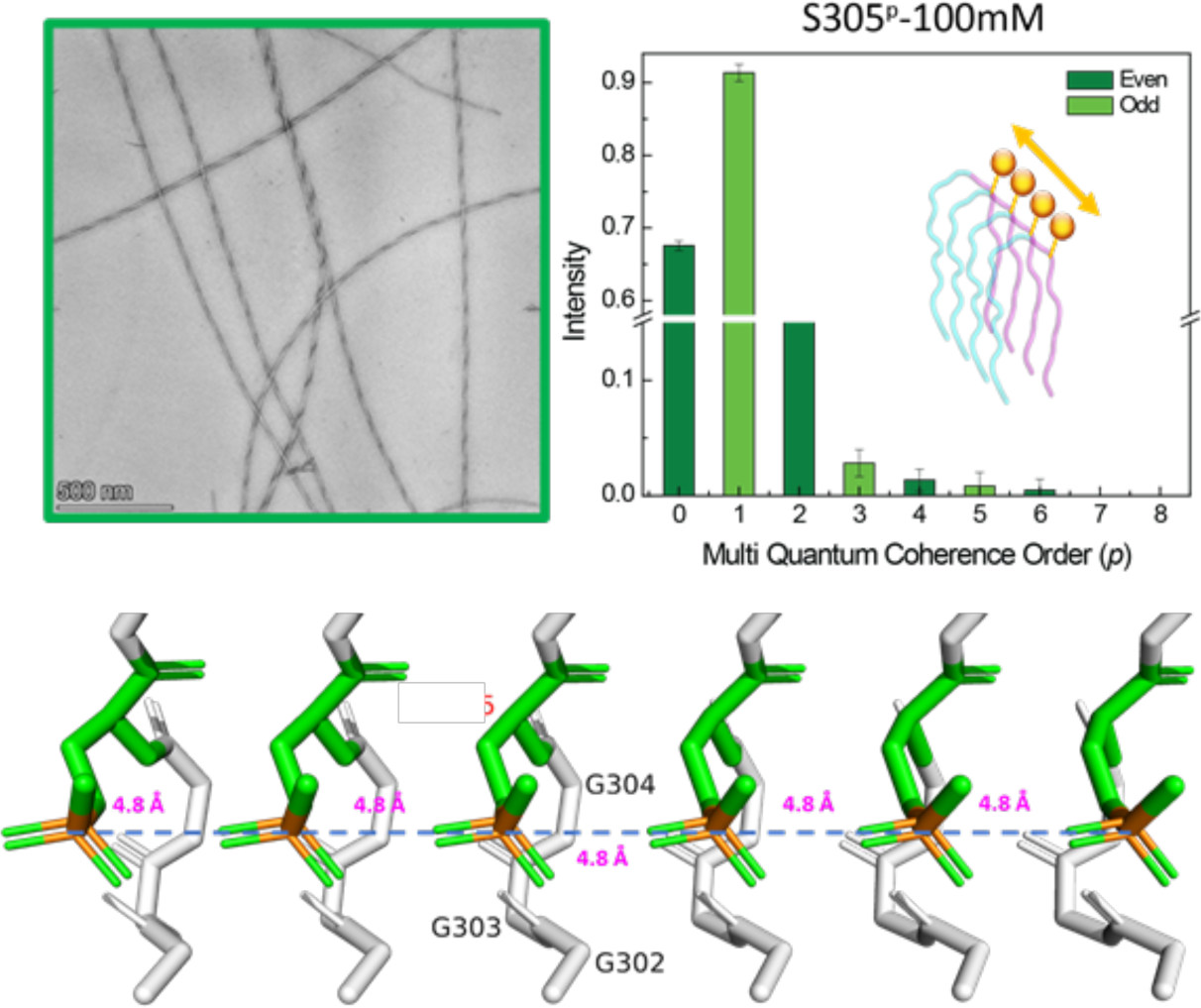

## Introduction

Tau is an intrinsically disordered protein (IDP) that serves many biological functions, most notably as a scaffold for stabilizing microtubules in neurons. Under physiological conditions, hyperphosphorylation of tau is proposed to play an essential role in regulating tau function and/or a hallmark of pathogenic condition.^1,2^ While the human tau protein isoforms exhibit phosphorylation at up to 40-50 out of 85 sites of serine, threonine, and tyrosine residues under pathological conditions^3^, the PTM combinations that co-occur are not currently known. Hyperphosphorylated tau proteins are key constituents of neurofibrillary tangles (NFTs)^4^ found under several neurodegenerative disease conditions referred to as tauopathies, including Alzheimer’s Disease (AD)^5–8^, Chronic Traumatic Encephalopathy (CTE)^9–11^, Pick’s Disease (PiD), Corticobasal Degeneration (CBD) ^12^, Progressive Supranuclear Palsy (PSP)^13^, and Argyrophilic Grain Disease (AGD)^14,15^. While the current understanding in biology and neuropathology is that the hyperphosphorylated tau and their fibrillar assemblies are the neurotoxic form of tau, it remains unclear whether hyperphosphorylation is an aggravating factor in, or simply a reflection and consequence of, the process of tau self-assembly into filaments.^16^ Research suggests that phosphorylation occurs before aggregation^3^, and phosphorylation at different sites has diverse impacts on biological processes and pathogenic developments.^4,16–18^ Hyperphosphorylation of soluble tau may contribute to neurodegeneration by weakening association of tau with microtubules, and either causing microtubules to destabilize or increase free intracellular tau available for aggregation.^19,20^ While there are downstream biological consequences of phosphorylation, this study addresses an important unknown in the field, namely whether there are direct, stand-alone, molecular consequences of phosphorylation on filament formation and stability *in vitro*, outside the cellular environment. This understanding would be key to the synthetic replication of tau NFTs found in tauopathies with molecularly defined protein folds and quaternary structures that are diagnostic hallmarks of different tauopathies including AD, CTE, CBD, PSP, AGD and several other diseases.

A recent discovery revealed that phosphate-containing molecules, including dissolved monophosphates and ATP, form dynamic assemblies under biological solution conditions in water.^21^ This hinted at the possibility of a hitherto unknown role of phosphorylation to promote self-assembly and stabilization of tau fibrils. The hydrogen bonding energy of phosphates may or may not be sufficiently strong to be a driving force for aggregation. However, once tau fibrils are formed, phosphoryl groups could be a major stabilizing factor for fibrils, given that phosphates readily form extended hydrogen-bonded networks. At the minimum, adjacent phosphoryl groups along the fibril’s elongating axis may not be a destabilizing factor due to its net negative charge that would be repulsive for closely spaced phosphoryl groups within 4.8 Å in β sheet arrangements.^22^ Experimental investigations are needed to test whether phosphate groups on tau proteins associate into a 1D network with long-range order, and whether they can act as stabilizing factors for tau filaments.

Cryo-EM has been transformative in elucidating the structure of tau fibrils from the postmortem brain of tauopathy patients^23–26^, while solution and solid-state NMR spectroscopy elucidated valuable structural information on pathological tau proteins^27–29^. Tau is hyperphosphorylated at several sites (e.g., S202, S262, S356, and S396) under tauopathy conditions. However, Cryo-EM structures of tau fibrils do not reveal atomic-level details about the occurrence and organization of phosphorylated groups, despite antibodies indicating their presence.^23,30,31^ This could be due to the heterogeneous nature of phosphorylated species or because many of these modifications are outside the fibril core. Research into the structural effects of phosphomimic mutations has shown that the S356D mutation impedes the formation of Alzheimer’s disease (AD)-like folds, whereas a combination of S396D, S400D, T403D, and S404D mutations promotes the formation of an AD- fibril core.^26^ Even so, the effect of pseudo-phosphorylation and actual phosphorylation would be distinct if the phosphoryl groups form an extended hydrogen-bonded network mediated by water. A recent study found that phosphorylated 2N4R tau with the enzyme GSK3β shows enhanced liquid-liquid phase separation (LLPS) and forms a fibril core with a C-shape fold similar to that found in AD.^16^ Solution state NMR reported on the effect of phosphorylation on both the transient local secondary structures and global rearrangement of the dynamic ensemble of modified tau.^27–31^ Certain combinations of phosphorylation, especially in the polyproline rich region (PRR), have been found to induce secondary structural changes.^32,33^ Phosphorylation of S262 in the R1 repeat domain has been shown to enhance the assembly rate of tau, while other studies suggest that the same modification hindered aggregation.^34–36^ Recent studies using NMR and paramagnetic relaxation interference (PRI) showed that phosphorylation at specific residues e.g. S202 and T205 within the PRR lead to confirmational changes in the tau ensemble, and presumably also the tau aggregation pathway.^37,38^ Solid-state NMR (SSNMR) and cryo-EM combined could determine the rigid core structure and the fuzzy coat property of phosphomimetic tau fibrils with S202E, T205E, and S208E modifications in AT8-3E tau and 396E, S400E, T403E, and S404E in PHF1-4E tau.^39^ However, none of these studies investigated the impact of phosphorylation on intermolecular interaction.

This study aims to analyze the spatial organization of phosphoryl groups in tau fibrils using SSNMR that captures through-space ^31^P-^31^P dipolar couplings to determine whether long-range arrangements of phosphoryl groups can be found. The only spectroscopic method that can offer such information is through the determination of coherence order by multiple quantum (MQ) spin counting (SC) SSNMR.^31^P MQ ssNMR has been used to establish the connectivity between ^31^P nuclei involved in the coupling network of solid-state phosphate clusters and crystals.^40,41^ Tycko’s group applied the ^13^C MQ-SC technique on solid-state powder samples of amyloid fibrils made of a seven residue Aβ16-22 fragment and full length Aβ1-40 to probe the supramolecular organization of the beta-sheets, and observed a MQ-CO up to 4 that suggests in-register, parallel beta-sheet stacking.^42,43^ However, these MQ-SC studies were of powdered solid state samples. We recently demonstrated that ^31^P MQ-SC can be applied to small clusters and molecules in solution state upon vitrification, which can capture species formed under native conditions, when combined with NMR signal enhancement by dynamic nuclear polarization (DNP).^44^ In the present work, we utilized the ^31^P MQ-SC technique in combination with DNP to investigate whether phosphoryl groups organize into extended networks in fibrils of phosphorylated tau fragments.

We use a critical tau fragment consisting of 19 residues spanning D295 to V313 of 4R tau, that forms the junction between the R2 and R3 repeat domains and contains the P301L mutation (jR2R3-P301L). This peptide forms tau fibrils containing pathological folds and display prion properties.^45,46^ Two sites within jR2R3-P301L may be phosphorylated under pathological conditions: S305 and Y310. We hence prepared phosphorylated (not pseudo-phosphorylated) jR2R3-P301L at either the S305 or Y310 sites, referred to as S305^p^ and Y310^p^, respectively. The study commenced by assessing the quality and stability of the phosphorylated jR2R3-P301L model tau peptide fibrils through thioflavin T fluorescence (ThT) assays and negative-stain transmission electron microscopy (TEM) imaging, followed by examining fibril aggregation by ThT. We then proceeded with ^31^P ssNMR MQ-SC studies to investigate whether the phosphoryl groups on specific sites of the peptide form a higher order, water-mediated, hydrogen-bonded network. We conducted numerical simulations on spin systems with various geometries to analyze the experimental ^31^P MQ-SC results, and gained insights into the minimal number of defect free in-register tau packing in the fibril or the formation of prototype paired helical filament geometry, while relying on the solved tau fibril structures by cryo-EM.^45^

## Results and discussion

### Aggregation assays

Phosphorylated jR2R3-P301L tau fibrils at serine 305 or tyrosine 310 were synthesized using the preparation method described in the Materials and Methods section. Figure 1 depicts the 19- residue jR2R3 peptide sequence (D295 – V313) derived from the R2 and R3 regions of the 2N4R tau isoform, and the phosphorylation mutations at S305 and Y310 illustrated in a CBD conformation^47^ with the monomer forms stacked in-register and parallel along the fibril axis. The S305^p^ and Y310^p^ fibrils were prepared in a solution of either 100 mM or 1M NaCl during aggregation. High salt concentration such as 1M NaCl screens electrostatic interactions and sequester water away from the protein, thereby amplifying the effect of the hydrophobic association of tau. The high salt concentration is not physiologically relevant, however, the influence of such conditions on the proteins of strong electrostatic screening and enhanced hydrophobic effects can readily represent a physiologically relevant condition.^48^ Hence, the high salt concentration of 1M NaCl was examined in addition to 100 mM NaCl to prepare the phosphorylated fibrils. A ThT assay was used to monitor the formation and stability of fibrils. The aggregation of S305^p^ and Y310^p^ fibrils exhibited strong fluorescence owing to binding of ThT (Figure 2), indicative of β-sheet formation by fibrillization. In particular, fibrils made of S305^p^ showed higher fluorescence than Y310^p^ prepared under either salt concentration. The morphology and final fibril formation of S305^p^ and Y310^p^ were examined using negative stain transmission electron microscopy (TEM). TEM images showed that S305^p^ exhibited higher quality fibrils displaying paired helical filaments and ribbon-like structures compared to Y310^p^ fibrils (shown in Figure 2).

**Figure 1.**
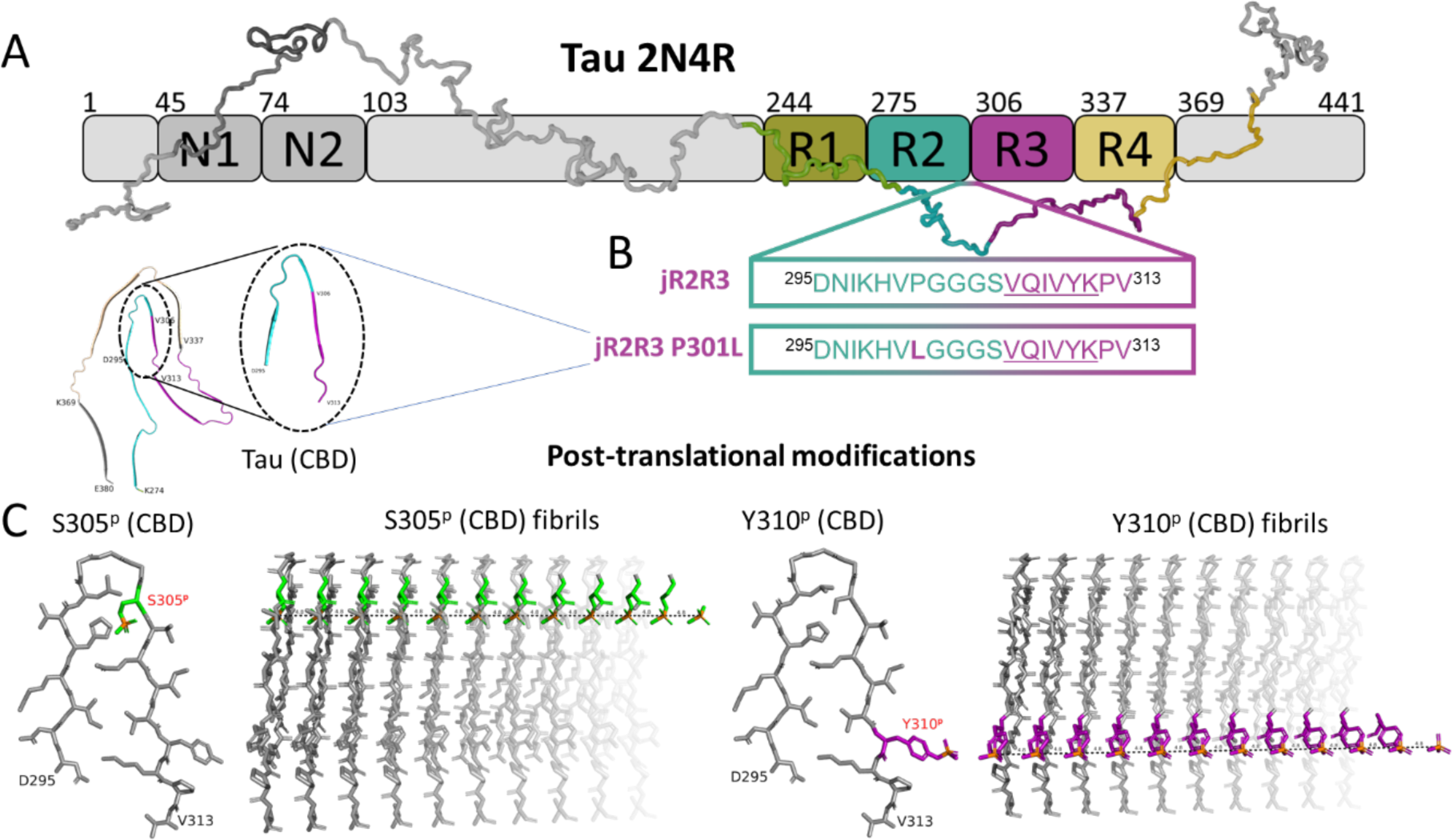
A) Schematic showing overview of the tau construct 2N4R. The elements are color-coded for clarity with R1 highlighted in Olive, R2 in Teal, R3 in Purple, and R4 in Sand. PDB structure representing 4R tauopathies CBD with PDB ID: 6VH7^45^ B) Strand-loop-strand designated as jR2R3 and jR2R3 P301L, are displayed with the PHF6 sequence underlined, and residues mutated relative to jR2R3 are shown in bold. jR2R3/jR2R3 P301L sequences represent the R2/R3 splice junction of tau spanning from D295 to V313. C) Post-translational modifications (PTMs) of jR2R3 P301L in CBD configuration are presented here. Serine 305 (Green) and Tyrosine 310 (Purple) are the residues at which the phosphoryl group is added. Stacking of the monomers is proposed as displayed in the above figure.

**Figure 2.**
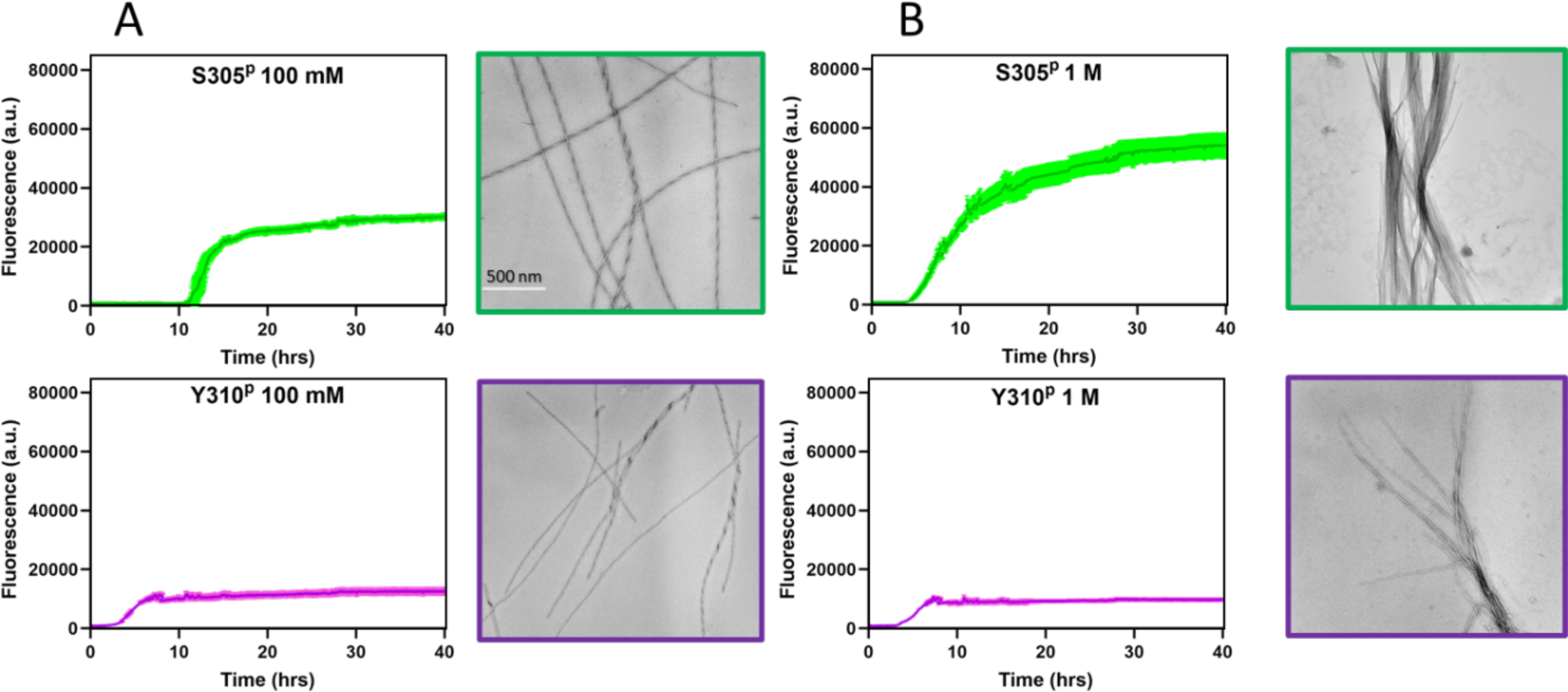
Monitoring the fibril aggregation of S305^p^ (Green) and Y310^p^ (Purple) jR2R3 P301L fibrils using the Thioflavin T (ThT) fluorescence assays and TEM micrographs for A) 100 mM NaCl and B) 1M NaCl conditions respectively. The scale bar is uniform for all TEM images.

Stability revealed by fiber denaturation experiments:

The stability of each the fibrils was tested by incubating the fibrils to guanidinium hydrochloride (GdnHCl) at varying concentration from 0 to 1 M. GdnHCl is a denaturant that can disrupt the natural structure of proteins by breaking hydrogen bonds and disturbing β-sheet secondary structures.^49^ The ThT fluorescence was measured after equilibrating the samples for 24 hours. Fibrils that withstand denaturation under more severe conditions are considered more stable. Therefore, a larger decrease in ThT fluorescence as GdnHCl concentration increases indicates lower fibril stability.

The stability of the jR2R3 P301L fibrils prepared under 100 mM NaCl condition are shown in Figures 3A and S6. To standardize the results, all values were compared to the initial fluorescence of the corresponding fibrils before denaturation. Under denaturing conditions with GdnHCl, all peptide fibrils exhibited reduced fluorescence intensity, as the fibers treated with GdnHCl at least partially disassemble into monomers. Notably, at 1M GdnHCl concentration, the peptide fibrils displayed distinct differences in their stability: S305^p^ fibrils had a net loss of 28% ± 0.96% and Y310^p^ a net loss of 52% ± 0.57% in fluorescence, whereas jR2R3 P301L fibrils showed a greater net loss of 63% ± 0.97%. The TEM images (Figure S6) exhibit the characteristic features of denaturation, with more fragmentation observed in Y310^p^ compared to S305^p^ or jR2R3 P301L fibrils with increasing GdnHCl concentration. These results show that phosphorylated jR2R3 fibrils are more stable compared to non-phosphorylated jR2R3 fibrils. Of these, S305^p^ fibrils exhibited the highest stability. These results suggest higher orders of associations in phosphorylated fibril. This hypothesis will be further examined in the upcoming sections.

**Figure 3.**
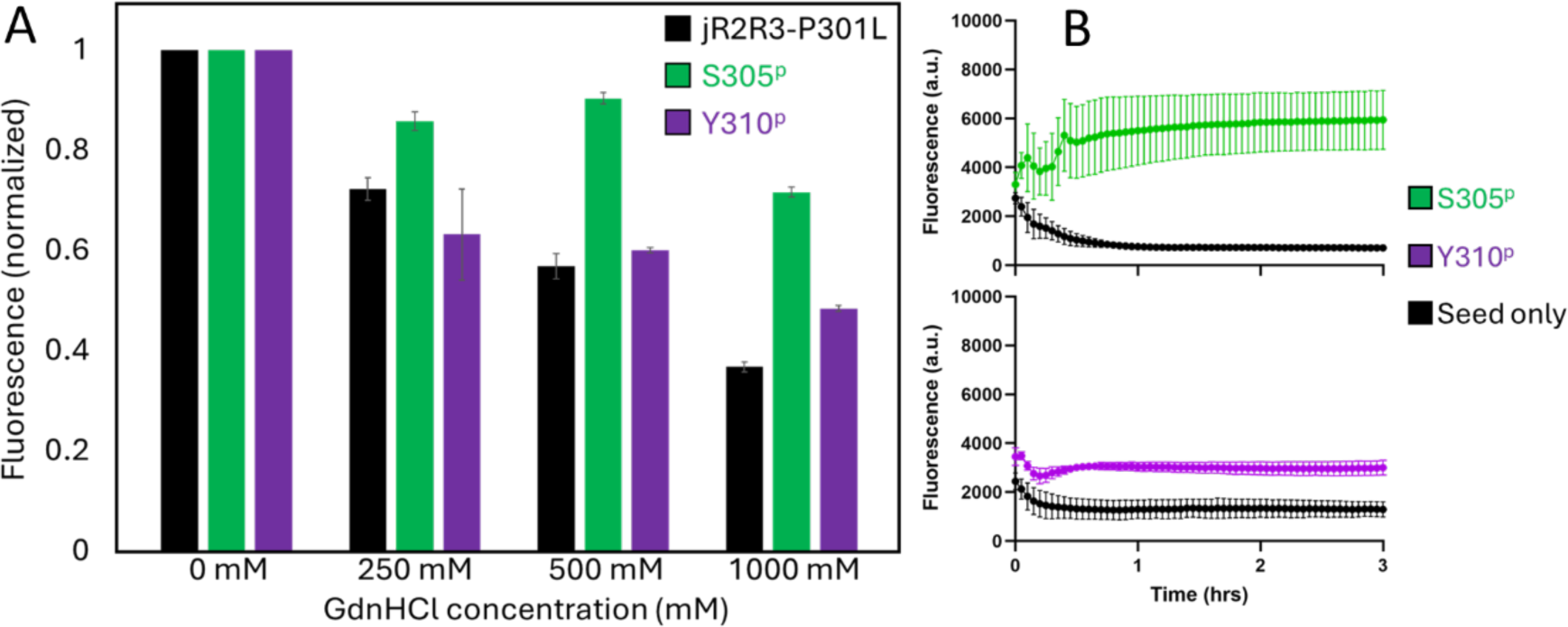
A) Fiber denaturation experiment on fibrils with 100 mM NaCl condition, normalized ThT fluorescence as a function of Guanidinium hydrochloride (GdnHCl) concentration after 24 hours of incubation at a temperature of 37 °C. S305^p^ retains the most fluorescence at the highest concentration, indicating a more stable fiber compared to both jR2R3-P301L and Y310^p^. B) Seeding, equal amounts of each seed (black) were added to their corresponding monomer type, S305^p^ shows the highest increase in ThT fluorescence, indicating more fiber formation compared to Y310^p^.

Seeding competency studies:

Since the S305^p^ fibrils are more stable compared to the Y310^p^ fibrils, we hypothesize that this would improve the fibers’ ability to be more seeding active. The templating of tau to adopt the same structural folding is a key attribute of a competent seed, and therefore a well-packed fiber should be properly aligned to allow additional monomers to stack on more readily upon adopting the fold of the fibrillar seed. Using absolute ThT fluorescence as a guide to assess seed performance, the comparison of the ThT fluorescence of the seed to the seeded monomer, we demonstrate that the S305^p^ fibrils are more seeding active than the Y310^p^ fibrils (Figure 3B). After analyzing the fibril morphology, stability, and quality, the S305^p^ and Y310^p^ fibril samples were scaled up for solid-state NMR studies.

### MAS DNP studies

Conventional one-dimensional ^1^H-^31^P CP spectra were recorded under MAS at 10 kHz and DNP at 100 K. The ^31^P NMR lineshape and width reflect the spatial distribution of phosphoryl groups. The ^31^P NMR spectra were analyzed using a Gaussian/Lorentzian model with the DMfit software (shown in Figure 4).^50^ The ^31^P NMR spectra of vitrified monomers of both S305^p^ and Y310^p^ show a broad Gaussian lineshape, as expected (Fig. 4A and 4D). The ^31^P NMR spectra of the S305^p^ fibrils prepared in 100 mM or 1M solution show a significantly narrowed line. Critically, the dominant component with a chemical shift of 0.99 ppm (70%) is a narrowed, Lorentzian line, with a full width at half maximum (FWHM) of 220 Hz compared to 800 Hz of the monomer sample with a Gaussian lineshape. The other two components of the S305^p^ fibril include one Lorentzian component centered at 4.4 ppm with an FWHM of 570 Hz (5 %) and a Gaussian component centered at -1.67 ppm with an FWHM of 1010 Hz (25 %). We attribute the two broader components to fibril populations with powder orientation of the ^31^P spectral components. In contrast, the ^31^P NMR spectra of the Y310^p^ fibrils, prepared in either 100 mM or 1M solution, showed no narrowed ^31^P NMR spectral component. The ^31^P NMR spectral narrowing, only seen in S305^p^ fibril, suggests that S305^p^ fibrils are stabilized by closely coupled phosphoryl groups. Random orientation of phosphoryl groups in space will result in a broadened Gaussian lineshape, as observed for Y310^p^ fibrils, while one-dimensionally stacked ^31^P would yield mutually commuting dipolar Hamiltonians that produce dipolarly-narrowed Lorentzian ^31^P NMR lineshapes, as observed in S305^p^ fibrils.

**Figure 4.**
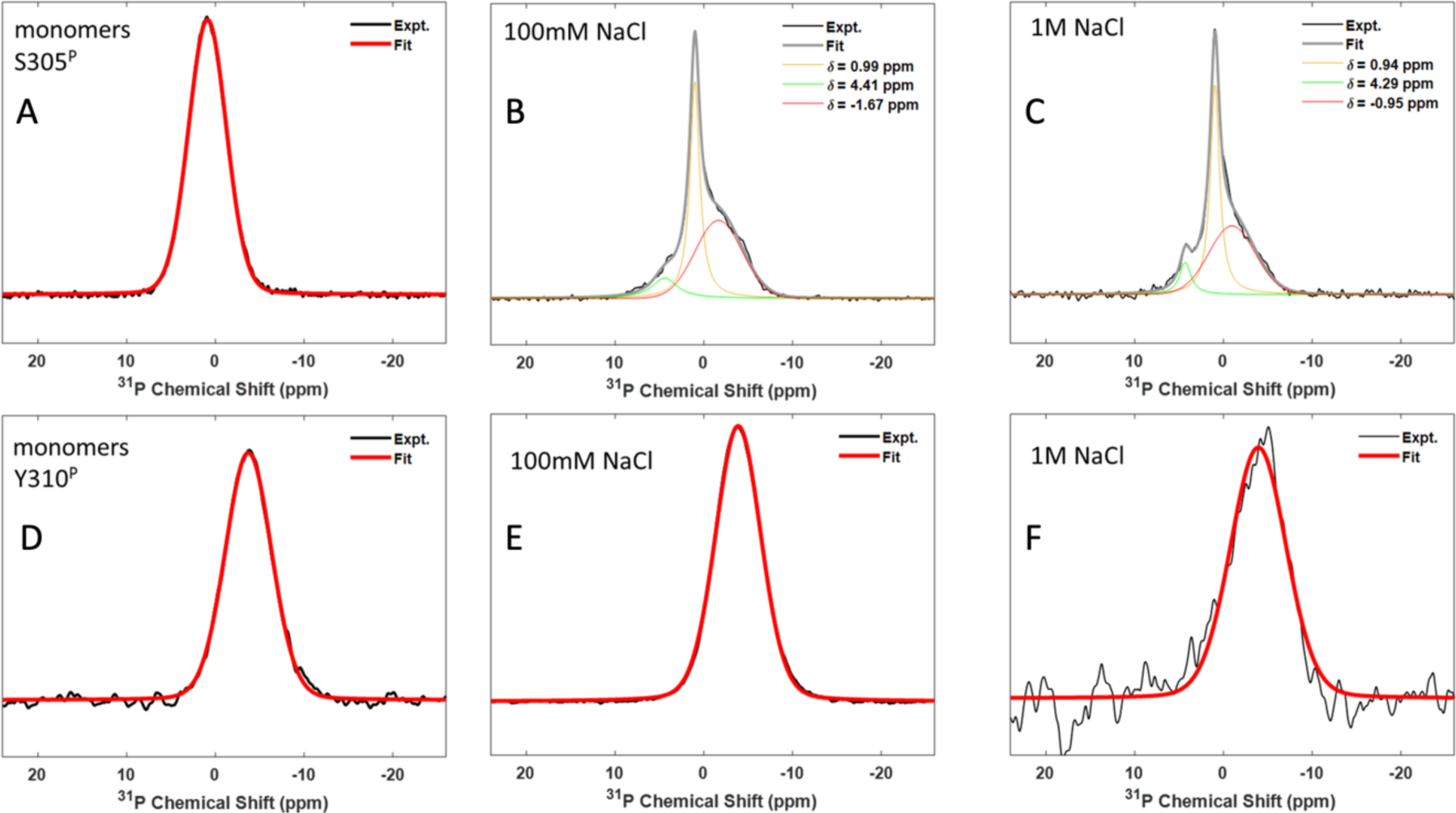
^31^P NMR spectra of S305^p^ jR2R3 P301L and Y310^p^ jR2R3 P301L samples and the corresponding deconvolution of the spectra using DMfit software. A) monomer, B) 1 M NaCl, and C) 100 mM NaCl condition of S305^p^ jR2R3 P301L samples, D) monomer, E) 1 M NaCl, and F) 100 mM NaCl condition of Y310^p^ jR2R3 P301L samples. Spectra were obtained at 10 kHz MAS frequency and 100 K temperature under DNP. A signal enhancement of 40-fold was attained with DNP for each spectrum. Each spectra was acquired with 128 scans for signal averaging except for 1M NaCl Y310^p^ and a recycle delay of 5 seconds was used. We applied coif3 wavelet denoising using NERD software from Cornell University to obtain a good signal-to-noise ratio for the 1M NaCl Y310^p^ spectra. A similar line broadening of 50 Hz was applied to each spectrum.

The line broadening in solid state NMR spectra is mainly caused by anisotropic interactions: chemical shift anisotropy (CSA), homonuclear, and heteronuclear dipolar couplings. CSA fits were performed using the CSA-MAS model in the DMfit^50^ software (fit values shown in Table S2). The CSA and asymmetry parameter (η) values for S305^p^ monomers and fibrils are similar, as well as for Y310^p^ fibrils and monomers, indicating CSA does not significantly contribute to the line width. The main sources of heteronuclear dipolar couplings are ^1^H-^31^P, ^13^C-^31^P, and ^15^N-^31^P dipolar couplings. Applying heteronuclear decoupling on protons during acquisition eliminated ^1^H-^31^P couplings. We confirmed this on the ^31^P NMR of 100 mM NaCl S305^p,^ showing a narrower and more intense peak at 0.99 ppm with decoupling on, while other peaks remained unaffected (Figure S2). The contribution from ^13^C-^31^P and ^15^N-^31^P couplings is negligible due to their smaller magnitudes at isotopic abundance, indicating heteronuclear dipolar couplings are not the primary source of spectral broadening. Therefore, the primary cause of line broadening is the ^31^P-^31^P homonuclear dipolar couplings. The reduction in linewidth observed in the ^31^P spectra of S305^p^ fibrils at both NaCl conditions, as depicted in Figure 4, indicates an averaging effect on the ^31^P- ^31^P homonuclear dipolar couplings. This averaging of the ^31^P homonuclear dipolar couplings can only be achieved by parallel stacking of the fibrils in a single orientation across several stacking copies of tau under MAS. In contrast, the ^31^P NMR spectra of Y310^p^ exhibited broad linewidths, similar to that of the monomer. This observation suggests that the incomplete averaging of homonuclear dipolar couplings in Y310^p^ fibrils is a result of heterogeneity or local disorder present within the sample contributing to the broad linewidths. Therefore, we can conclude that S305^p^ fibrils are lined by one-dimensional (1D) phosphoryl wires with better local homogeneity/order compared to Y310^p^ fibrils.

### MQ-SC experiments

We next carried out MQ-SC experiments to obtain direct evidence for phosphoryl group clustering and 1D ordering. Our goal is to determine the minimal cluster size by analyzing the highest possible extracted MQ coherence order, which is equal to the number of coupled spins present within a spin cluster. In the structure of the phosphorylated jR2R3 P301L fibrils, the positions of ^31^P spins in each S305^p^ or Y310^p^ residue of peptide molecule provide valuable insights into the supramolecular organization within the β-sheets, which lead to discussion on different spin network geometries such as parallel, anti-parallel, dimers, trimers, etc. with an internuclear distance of 4.8 Å.^22^ In an in-register, parallel β-sheet structure, the ^31^P spins on neighboring tau fibrils form a nearly linear arrangement with internuclear distances of approximately 4.8 Å.

MQ-SC experiments were performed on both fibril and monomer samples of S305^p^ and Y310^p^ at both 100 mM and 1M NaCl concentrations. We recently demonstrated the utility of the MQ-SC method on vitrified samples of adenosine triphosphate (ATP), amorphous calcium phosphate (ACP), crystalline hydroxyapatite (HAp) (Ca5(PO4)3OH, and simulated body fluid (mSBF) under DNP using SR2^1^8 pulse sequence to determine the number of ^31^P spins present in the clusters.^44^ The SR2^1^8 sequence is an R-symmetry-based pulse sequence^44,51,52^ used for double quantum excitation and reconversion in the MQ-SC experiments, and is a super cycled version of R2^1^8, described as a π/2-pulse sandwiched R2^1^8R2^−1^8 pulse sequence that is minimally sensitive to CSA. It requires the radio frequency power to be set to equal to half the MAS frequency. The excitation time of 8 ms was found to be optimal when the MQ-SC experiment was performed on 1M NaCl S305^p^ fibrils with different excitation times of SR2^1^8 sequence (shown in Figure S3). The same excitation time of 8 ms was used during MQ-SC experiments on all fibrils/monomers in this study. The phase of the excitation block is incremented in a phase step 2π/j (j=number of experiments) to extract the multi-quantum coherences, and all the remaining phases in the experiment are kept constant. Figure 5 illustrates the MQ-SC profiles of even and odd spin-counting and the corresponding coherence orders extracted for S305^p^ and Y310^p^ respectively for 100 mM and 1M NaCl conditions. Here, we utilized multi-cosine fitting to extract the MQCOs, as demonstrated in a recent study of our group^44^, that relies on the knowledge that the spin counting profile takes the form of multi-cosine waveforms as elucidated by the Average Hamiltonian Theory described by Olyer and Tycko^53^. The spectra of S305^p^ fibrils prepared at 100 mM or 1M NaCl concentration showed the highest (MQCOs) up to four. In contrast, the maximum MQCO observed for Y310^p^ fibrils was two, which can originate from a pair of dipolar coupled phosphoryl groups, suggesting an absence of in-register stacking of phosphates in Y310^p^ fibrils.

**Figure 5.**
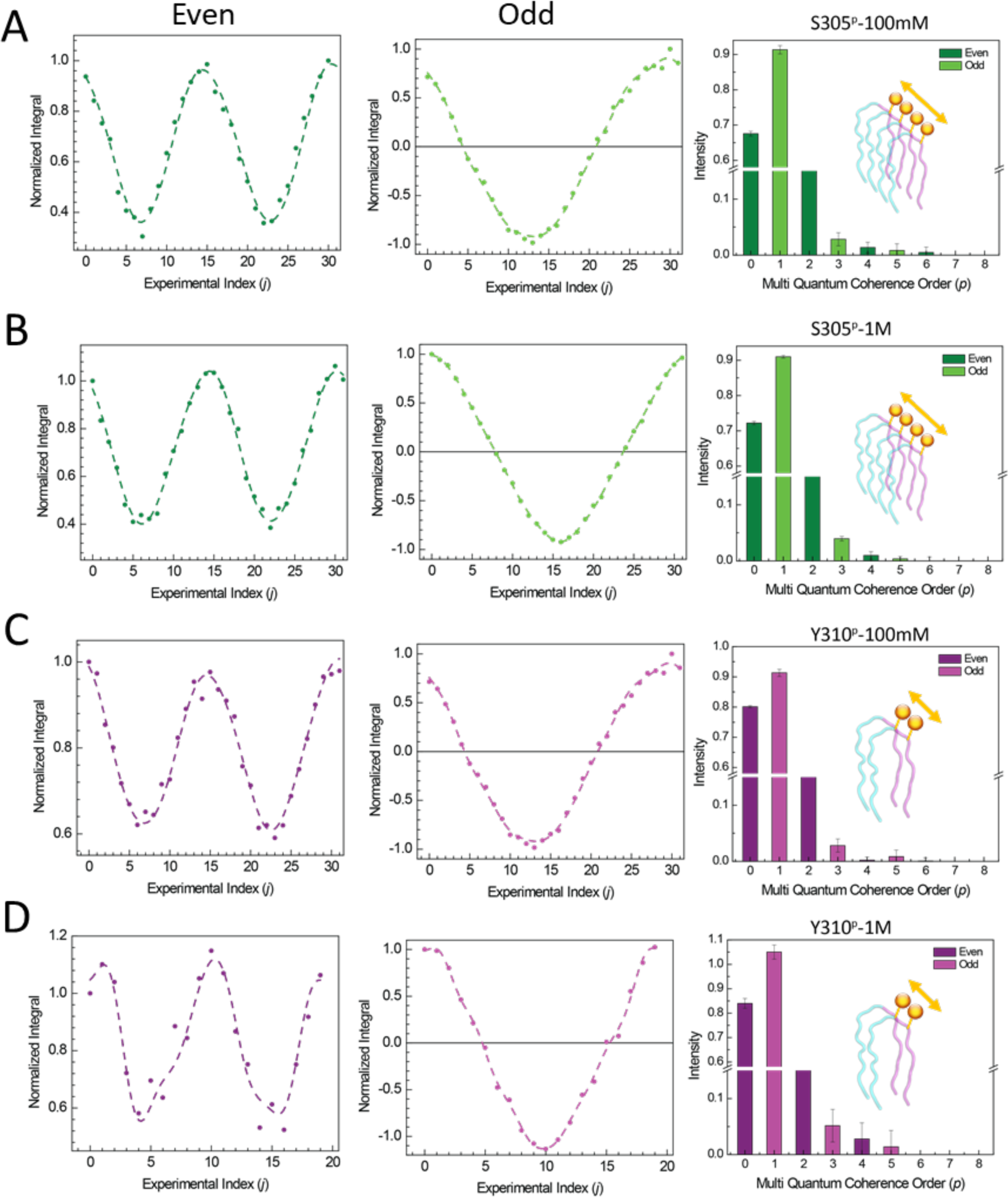
Even and odd spin counting profiles and MǪCO profiles obtained from multi-cosine function curve fitting of S305^p^ jR2R3 P301L fibrils and Y310^p^ jR2R3 P301L fibrils at 10 kHz MAS frequency and with 8 ms excitation time during SR2^1^ DQ recoupling sequence under DNP and a repetition time of 5 s. (A) 100 mM NaCl S305^p^, (B) 1 M NaCl S305^p^, (C) 100 mM NaCl Y310^p^, and (D) 1 M NaCl Y310^p^. The x-axis of the spin counting profiles is represented by the experimental index (j), where each phase is incremented by 360°/experimental index. All spin-counting profiles’ integrals were normalized to the integral of the first experiment (j = 0).

To investigate the potential structure of S305^p^ fibrils that give rise to an MQCO of four, we performed MQ-SC simulations with six ^31^P geometry networks using the SIMPSON software.^54–56^ The SIMPSON simulations of MQ-SC were conducted using parameters identical to those employed in the experimental setup, with the spin systems of different geometries described in Table S3. Even and odd MQ-SC simulations were carried out for different geometries, such as a dimer, trimer, linear chain of six spins, and prototypical paired helical filament (PHF) geometry. This allowed for an exploration of the spin arrangements and their corresponding multi-quantum coherences in each of these geometries. A distance of 4.8 Å was used between two successive ^31^P spins, corresponding to the inter-strand β-sheet distance.^22^ Figure 6 presents the results of the MQ- SC simulations that show the extracted MQCOs for different geometries. Dimer geometry simulations (Figure 6a) produced MQCOs of 1 and 2 with odd and even spin counting simulations, respectively, as expected from the spin system. The MQSC simulations of the trimer spin system showed the MQCOs of 1, 3, and 2 for odd and even spin counting simulations, respectively, according to the expected MQCOs of 1, 2, and 3 in the trimer spin system. Six spin network simulations of the linear arrangement of ^31^P spins showed the MQCOs of 1 and 3 with odd spin counting and MQCOs of 2 and 4 with even spin counting, respectively, displayed in Figure 6c, not exceeding the MQCO of 4 above the noise level in the MQCO charts. Spin counting simulations for a paired helical geometry displayed MQCOs of 2 and 4 with even spin counting simulations and 1 and 3 with odd spin counting simulations, respectively. Based on the comparison of simulation results of the linear chain comprising six spins and experimental results of S305^p^, it indicates that ^31^P spins in each peptide molecule in the fibril are adopting a parallel in-register packing arrangement of the beta sheets. In addition, the simulations of the prototypical PHF geometry are also plausible structures, even though these structures lead to higher MQCOs than measured. With the decreasing probability of observing higher-order quantum coherence and the stringent requirement of T1 and T2 relaxation of ^31^P NMR, weak dipolar couplings between ^31^P spins as well as isotropy of ^31^P chemical shifts to successfully carry out MQ-SC experiments, it is exceedingly difficult to observe MQCOs exceeding four, regardless of the defect-free order number of phosphoryl groups. Therefore, a measured MQCO of four in fibrils made of phosphorylated S305 tau peptides implies that there is in-register parallel β-sheet stacking with neat phosphoryl group ordering, while the data is also consistent with prototypical PHF geometries. In contrast, Y310^P^ fibrils did not show MQCOs of more than two, indicating a lack of parallel in-register stacking. The MQ-SC experiments show that the phosphoryl groups form tightly ordered stacks that reinforce the supramolecular fibril assembly despite the net negative charge of the phosphoryl groups that were assumed to experience repulsive forces. Our observation suggests that water-mediated, H-bonded, clustering of phosphoryl groups is favorable, that in turn impacts the folding and aggregation pathway and resulting fibril property. The MQ-SC results are consistent with a 1D array of the phosphoryl groups in S305^p^ fibrils that lead to line narrowing of the ^31^P spectra, and random distribution of phosphoryl groups in Y310^p^ fibrils causing broad line widths. Control MQ-SC experiments were performed on monomeric S305^p^ and Y310^p^ peptides not subjected to aggregation. The MQ-SC profiles of the monomers show random fluctuations (displayed in Figure S5) as a function of phase increment of the excitation block, and accordingly no MQCO could be counted upon Fourier Transformation. This is expected since there is no local ordering in the monomer samples in the absence of fibrillization. We emphasize that MQ-SC is the only method to offer direct insight into the number of ordered phosphates in extended structures, sort of directly visualizing the spatial ordering of phosphoryl groups.

**Figure 6.**
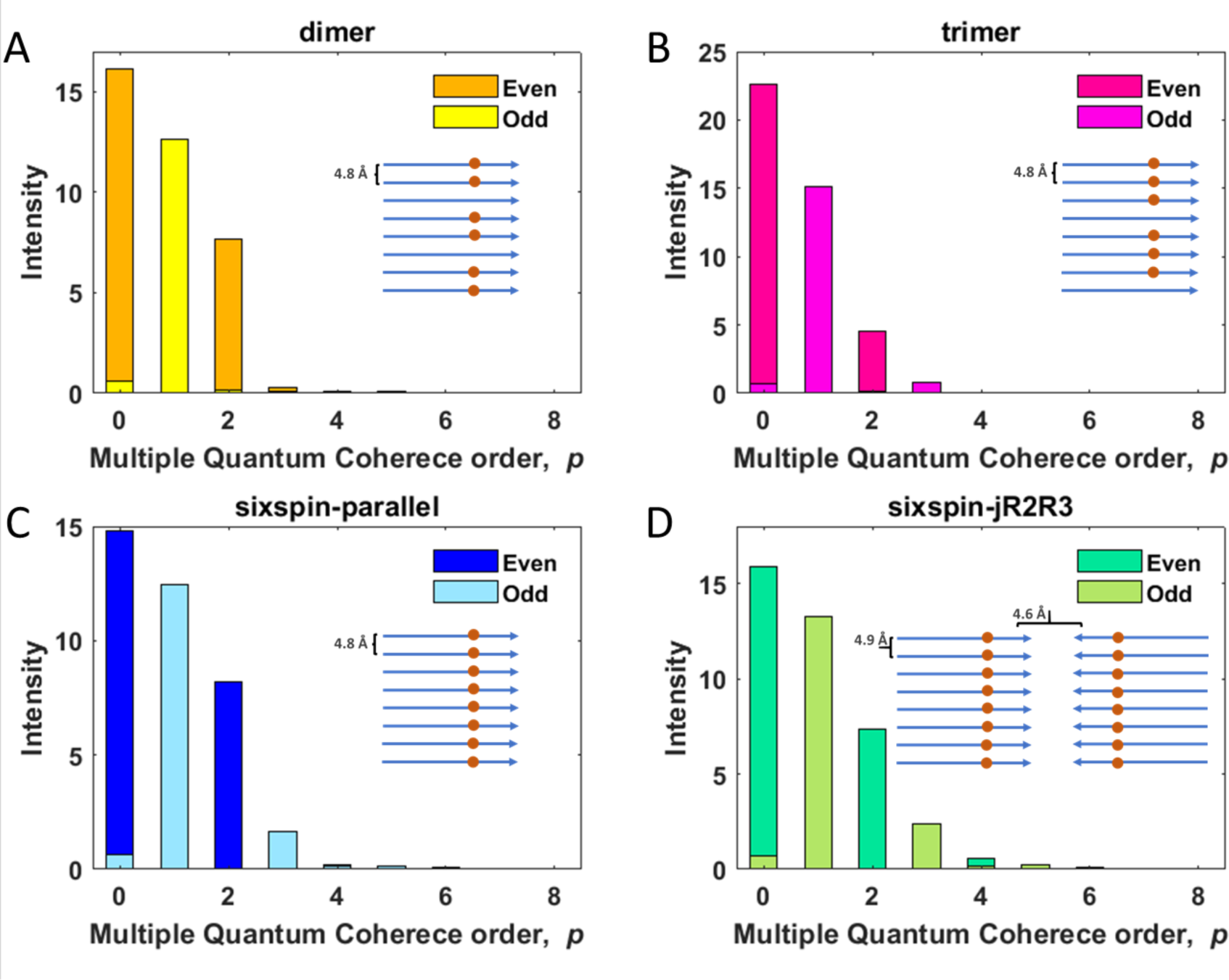
Numerical SIMPSON simulations of MQ-SC coherence orders with different geometries of a six-spin system: A) Dimer, B) Trimer, C) six-spin parallel configuration, and D) six-spin geometry with structure jR2R3 solved from our lab. Description of the spin system of each configuration is given in Table S3.

### Models/Hypothesis

Taken together, S305^p^ fibrils showed a MQCO of four and Y310^p^ fibrils of two to three, while the S305^p^ fibrils were significantly more stable under denaturation conditions. This indicates that the S305^p^ fibrils form higher order assemblies, and that the phosphoryl groups of site S305 tightly arrange by parallel, in register, stacking that hence stabilize the fibril integrity. In contrast, phosphorylation at Y310 does not give rise to an appreciable MQCO number. Phosphorylation at S305 may be favorable for stabilizing the CBD conformer by stabilizing a salt bridge with site H299, while phosphorylation at Y310 may be favorable for the PSP conformer by stabilizing a salt bridge with site H299. Our results show that phosphorylation at specific sites can favor or disrupt specific misfolding pathways depending on the favored folding pathway for the given aggregation condition. In simpler terms, phosphorylation can either bolster aggregation, seeding competency and fibril stability, or it can interfere with the pathological characteristics of tau fibrils. Thus, the effects of phosphorylation on individual sites are highly dependent on the specific location and dominant misfolding pathway. Hence, the outcomes of combinatorial phosphorylation should be highly context dependent. Nevertheless, we can deduce a novel insight into the role of phosphorylation in tau aggregation: phosphorylation can simultaneously assume the role of promoting or hindering intra-tau misfolding tendencies and will augment fibril stability if the phosphorylated site becomes part of a stable core within the tau fibril structure.

## Materials and methods

### Fibril preparation method

All jR2R3 P301L peptides were purchased from GenScript and powders were dissolved to 1 mM stock solutions and stored at -80° C or used immediately. The aggregation process was performed in a 50 mL Eppendorf tube. The stock peptide was diluted to 50 µM using 20 mM ammonium acetate, pH 7.4 (calibrated with HCl and NaOH), with either 100 mM or 1 M NaCl and accompanied by a 4:1 (peptide to heparin) molar ratio of additional heparin (average molecular weight 16 kDa). Continuous shaking was performed at 200 rpm at 37° C for 24 hours. Fibers were collected into a 50 kDa cutoff concentrator and rinsed 3 times with MilliQ water to remove monomeric peptide and excess heparin. Next, fibers were transferred to a weighed Eppendorf tube and lyophilized using a FreeZone 2.5 Liter -84° C Benchtop Freeze Dryer overnight. The mass of each fiber sample was quantified.

### Thioflavin T aggregation assay

All experiments were performed with a BioTek Synergy 2 fluorescent plate reader. In each well jR2R3 P301L (50 µM), ThT (20 µM), and heparin (12.5 µM) in buffer were distributed into a 384- well plate (Corning low volume non-binding surface black with clear flat bottom) to a total volume of 30 µL. The plate reader temperature was set to 37 °C and allowed to equilibrate, the samples were shaken under the programmed high shaking speed in between measurements. ThT fluorescence intensity was measured at (excitation=440 nm, emission=485 nm) every 2 minutes until a plateau was reached. These experiments were done in triplicate at least three times with independent samples.

### TEM analysis

For transmission electron microscopy (TEM) analysis, five μL of fibril samples were applied to a glow-discharged copper grid (Electron Microscopy Science, FCF-200-Cu) for 20 s and blotted dry with filter paper. Samples were stained with 5 μL 1.5 w/v % uranyl acetate and immediately blotted dry. An additional 5 µL of uranyl acetate was added for 60 seconds and blotted dry. Samples were analyzed using a Thermo Scientific Talos G2 200X TEM/STEM microscope operated at 200 kV and room temperature. Grids were then imaged with a Ceta II CMOS 4k x 4k camera at the indicated magnifications.

### Fibril stability by denaturation

Guanidinium Hydrochloride denaturation assays were performed using identical conditions to the initial fibrilization ThT assays. Prior to the stability tests and to ensure accurate ThT assessment during stability testing, it was necessary to remove any excess monomers and heparin. This was achieved by washing the fibrils in their original buffer containing 20 mM ammonium acetate and 100 mM NaCl using a concentrator with a 50 kDa cutoff. To achieve the specified GdnHCl concentration for the reported results, the fibrils’ initial concentration was reduced through dilution into the denaturation buffer. The values reported were obtained following 24 hours after the signal reached equilibration upon initial incubation.

### In vitro seeded aggregation protocols

Fibers used for the seeding experiments were assembled using identical conditions to the initial fibrilization assay. Samples were reconstituted to a final concentration of 1 mM in the reaction buffer. The seeds were subjected to sonication for 30 seconds using the microtip of a Qsonica sonicator, operating at a 30% duty cycle. Seeding assays were performed in a Tecan Spark, replicates were pipetted in a 384 well plate clear with black bottom, at a final concentration of 50 µM and a total volume of 30 µL. Seeds were added in a 1 monomer: 1 seed ratio.

### Solid-State NMR (SSNMR) experimental details

The lyophilized powdered fibrils were dissolved into DNP juice, prepared with 6:3:1 parts of glycerol-d8, D2O, and H2O with 10 mM AMUPol (Cortecnet, C010P005). The above samples with a final concentration of 4 mM (0.5 mg of fibril in 40 μl of DNP juice) were centrifuged into regular walled sapphire 3.2 mm MAS rotors and plugged with Teflon or silicon inserts to prevent leakage and subsequently injected into the spectrometer at 100 K. All the MAS DNP measurements were carried out on a Bruker Avance III DNP-NMR 400 MHz spectrometer equipped with a 25 W gyrotron microwave source (operating at 263 GHz frequency) at a temperature of 100 K. A triple resonance H/X/Y probe was used in double resonance mode with a 50 Ohm resistor on the Y channel. The signal enhancement of 40-fold was obtained using the DNP method with samples in the sub milligrams (0.4 to 0.5 mg) regime. One-dimensional ^31^P spectra were collected by using ^1^H-^31^PC cross-polarization experiments. MQ-SC experimental data was obtained using the SR2^1^8 pulse sequence for DQ excitation and reconversion. Further experimental details were given in the SI.

### Quantum mechanical simulations of SSNMR experiments

Multi-quantum SSNMR simulations were carried out using the SIMPSON package with a magnetic field strength of 9.4 T and MAS frequency of 10 kHz. SR2^1^8 pulse sequence was used for excitation and the reconversion blocks with a radiofrequency amplitude of 5 kHz, equal to half the spinning frequency which is the requirement for fulfilling the basic structure of the R2^1^8 symmetry sequence. 8 ms time was used for the excitation and reconversion blocks. Powder averaging was performed with a crystal file rep20 and ten gamma angles. Simulations for six ^31^P spin networks with different geometries were performed to explain and support the experimental results. The spin system was generated using the MagResView^57^ software for different geometries. Isotropic chemical shifts and scalar couplings were not considered in the simulations. The phase of the excitation block was incremented with a step of π/16 while keeping all other phases constant. The multi-quantum coherences were extracted by applying a Fourier Transform to the intensity profile obtained from the simulation.

## Conclusion

The molecular-level understanding of how PTMs, such as phosphorylation, influence tau aggregation is still lacking. In this regard, we investigated the spatial arrangements of the phosphoryl groups in tau fibrils in order to shed light on the stability and aggregation of the tau fibrils. The one-dimensional ^31^P spectra and MQ-COs revealed that S305^p^ fibrils had significant ^31^P line-width narrowing and can generate at least four ^31^P MQCOs, while Y310^p^ fibrils showed no line narrowing compared to the respective monomers and only up to two ^31^P coherence orders. This indicates that in jR2R3 P301L fibrils, phosphorylation at S305 enhances ordering and stability, while phosphorylation at Y310 does not. Based on these experimental findings, we present a new role of tau protein phosphorylation of enhancing tau aggregation propensity and fortifying the tau fibril stability by tight, in register, packing of phosphoryl groups in one-dimensional (1D) arrangements. We propose that the 1D phosphoryl wires are stabilized by water-mediated hydrogen bonding of the phosphoryl groups. Given that hyperphosphorylation is one of the most common PTMs and hence a biomarker of pathological tau proteins under neurodegenerative disease condition, we propose that fortifying tau fibrils by 1D phosphoryl wires is a relevant mechanism, depending on the specific combinatorics of the phosphoryl groups, that stabilizes pathological tau fibrils. It has been observed that more stable fibrils with greater ordering also display greater seeding competency. We hence conjecture that phosphoryl group fortified S305^p^ tau fibrils have enhanced toxicity given their greater seeding competency.

This study of a 19-residue peptides with only two potential phosphoryl sites demonstrates that the effects of phosphorylation of even a single site is highly site-specific and context dependent, and hence the effects of combinatorial phosphorylation are expected to be even more complex. Still, we learned that phosphorylation can play dual roles of enforcing or disrupting intra-tau misfolding and contribute to enhanced fibril stability if the phosphorylated site is included in a stable tau fibril core. The present study provides a new approach and concept to investigate and rationalize the phosphorylation effect at different sites of tau proteins on their aggregation pathway. The future application of MQ-SC to various forms of tau fibrils such as phosphorylated, unphosphorylated, and various mutations will continue to shed light on the aggregation pathways in tau pathologies.

## Acknowledgements

MSN thanks the National Science Foundation (NSF) Graduate Research Fellowship under Grant No. 1650114 and Heising-Simons Foundation for support. SH was supported by the NIH MIRA Award No. R35GM136411 for the fundamental study of phosphorylation and hydration and NIH NIA for the study of jR2R3 mini prion project. SH also thanks NIH Rainwater for the study of tauopathy-specific fibril formation. Electron microscopy was conducted at the Microscopy and Microanalysis facility at UCSB, a part of the materials research lab (MRL). This work made use of the MRL Shared Experimental Facilities supported by the MRSEC Program of the NSF under Grant DMR 1720256, a member of the NSF-funded Materials Research Facilities Network.

