## Supplementary file for "Phosphoryl group wires stabilize pathological tau fibrils as revealed by multiple quantum spin counting NMR"

Table of contents:

1. Extended methods

2. Overlay of $^{31}$P NMR spectra of S305$^{p}$ jR2R3 P301L and Y310$^{p}$ jR2R3 P301L samples

Table S1. Line shape fitting of the $^{31}$P spectra by Gaussian/Lorentzian using DMfit

Table S2. CSA parameters obtained from DMfit using the CSA-MAS model

3. Decoupling effect on $^{31}$P peak of 100mM S305$^{p}$.

4. Multi-quantum spin counting (MQSC) experiments with different SR$2_8^1$ mixing times on S305$^{p}$ SLS2 fibrils.

5. MQSC results on S305$^{p}$ and Y310$^{p}$ by conventional Fourier Transform (FT) method and also on monomers.

6. MQSC results on monomer samples.

Table S3. Description of the spin system used for SIMPSON simulations in Figure 6.

7. TEM images of stability tests on jR2R3 P301L, S305$^{p}$, and Y310$^{p}$.

1. Extended methods

NMR experimental details

1D $^{31}P$ CP spectra:

A pulse length of 2.8 μs was used for the excitation of the protons. For $^{31}P$, a pulse of length 3.4 μs was used for 90º rotation. $^{1}H$-$^{31}P$ cross-polarization experiments were performed with a ramp on the $^{1}H$ channel by varying its rf amplitude from 70 % to 100 % and 94 kHz rf power on $^{1}H$ and 80 kHz rf power on the $^{31}P$ channel, respectively with sufficient contact time. During the $^{31}P$ acquisition, heteronuclear decoupling of 90 kHz rf power was employed on the $^{1}H$ channel. One-dimensional $^{1}H$-$^{31}P$ CP spectra were obtained by 128 transients for each sample with a recycle delay of 5 seconds.

MQ-SC spectra:

MQ-SC experiments were carried out using the $SR2^{1}_{8}$ pulse sequence for DQ excitation and reconversion. $SR2^{1}_{8}$ works with DQ operators, and the spin counting experiments would generate an even number of multi-quantum coherences. The same $SR2^{1}_{8}$ sequence was used to get the odd number of multi-quantum coherences by shifting the starting coherence to ±1 with a 90º pulse applied before and after the DQ excitation and reconversion, respectively. During the $^{31}P$ acquisition and recoupling duration, heteronuclear decoupling of 90 kHz rf power was employed on the $^{1}H$ channel.  MQ-SC experiments were carried out with 256 scans. A relaxation delay of 5 seconds was used.

Data Processing

All NMR integrals taken for spin counting analysis were processed using a custom Python Jupyter Notebook code. Left shift, line broadening, baseline correction, and custom integrals were applied for all FIDs of a given data set similar to the procedure in our previous article.[1] MQCO profiles were very susceptible to phasing and left shift accuracy. Each FID was analyzed separately to ensure data was cohesive.

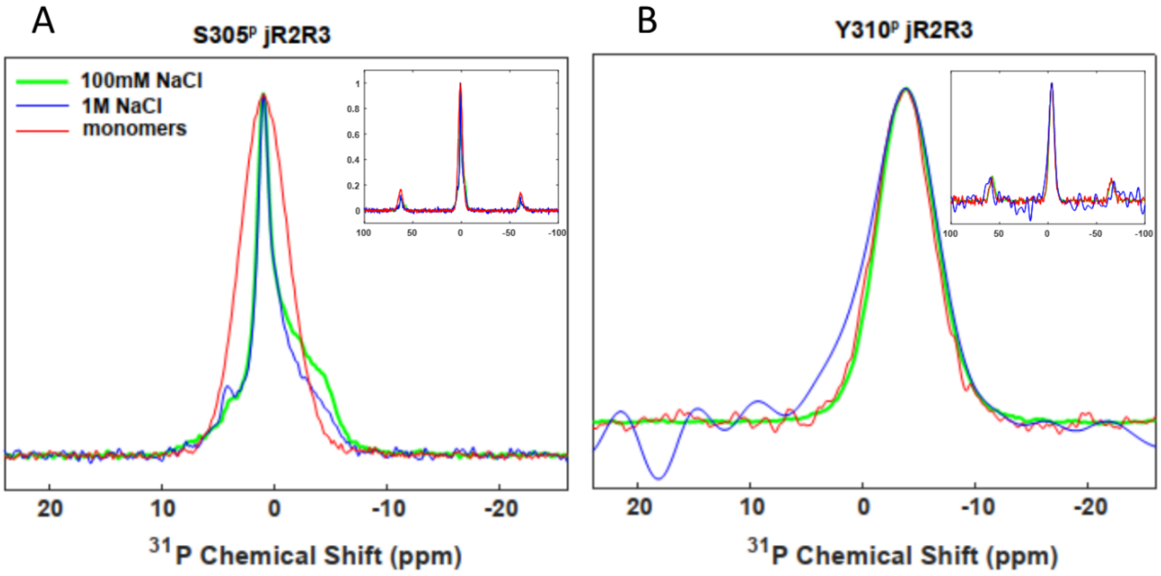

Figure S1. Overlay of $^{31}$P NMR spectra of A) S305$^{p}$ jR2R3 P301L and B) Y310$^{p}$ jR2R3 P301L samples, respectively.

Table S1. Line shape fitting of the $^{31}P$ spectra by Gaussian/Lorentzian using DMfit.

| **Sample type** | **0.9 ppm** | | **4.4 ppm** | | **-1.6 ppm** | |
|---|---|---|---|---|---|---|
| **S305$^{p}$** | **G** | **L** | **G** | **L** | **G** | **L** |
| 100mM_NaCl | 0.0 | 1.0 | 0.0 | 1.0 | 1.0 | 0.0 |
| 1M_NaCl | 0.0 | 1.0 | 0.0 | 1.0 | 0.95 | 0.05 |
| Monomers | 0.92 | 0.08 | -- | -- | -- | -- |
| 100mM_NaCl_nodec | 0.0 | 1.0 | 1.0 | 0.0 | 1.0 | 0.0 |
| **Y310$^{p}$** | **-3.9 ppm** | | | | | |
| 100mM_NaCl | 0.85 | 0.15 | | | | |
| 1M_NaCl | 0.9 | 0.1 | | | | |
| Monomers | 0.9 | 0.1 | | | | |

Table S2. CSA parameters obtained from DMfit using the CSA-MAS model

| Sample type | $\delta_{iso}$ (ppm) | CSA (ppm) | Asymmetry parameter (η) | Linewidth (Hz) |
|---|---|---|---|---|
| 100mM_NaCl_S305$^{p}$ | 0.99 | -72.0 | 0.5 | 300 |
| 1M_NaCl_S305$^{p}$ | 0.94 | -67.0 | 0.8 | 290 |
| Monomers_S305$^{p}$ | 0.95 | -85.0 | 0.75 | 800 |
| 100mM_NaCl_Y310$^{p}$ | -3.86 | -91 | 0.66 | 980 |
| 1M_NaCl_Y310$^{p}$ | -3.92 | -73 | 0.5 | 1000 |
| Monomers_Y310$^{p}$ | -3.81 | -81 | 0.95 | 960 |

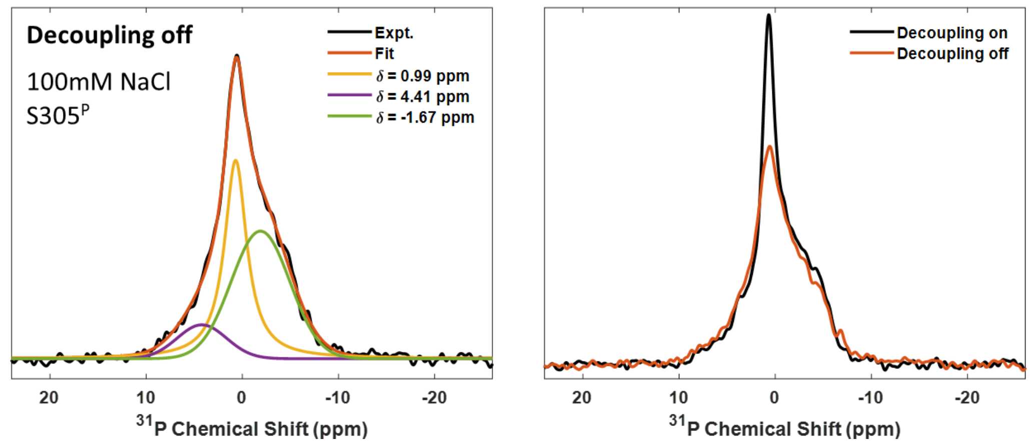

Figure S2. The effect of $^{1}H$-$^{31}P$ heteronuclear decoupling on $^{31}P$ spectra of 100 mM NaCl S305$^{p}$ jR2R3 P301L fibrils using the decoupling power on (black) and power off (red) on proton channel.

Decoupling effect on the $^{31}P$ spectra:

We verified the effect of heteronuclear decoupling on $^{31}P$ spectra of 100 mM NaCl S305$^{p}$ using the decoupling power on and power off (shown in Figure S2) on the proton channel during the acquisition. With decoupling power applied on protons, the peak at 0.99 ppm becomes much narrower with an increase in intensity when compared to the spectra with the power-off condition. The rest of the peaks were not much affected. We deconvoluted the $^{31}P$ spectra obtained by decoupling off condition and shown in Figure S2A. The peak at 0.99 ppm exhibited the Lorentzian line shape in both cases when the decoupling was applied on $^{1}H$ (Figure S1A) and not applied on. However, the peak at 4.4 ppm showed a Lorentzian nature when the decoupling was applied on $^{1}H$ and showed a Gaussian nature when the decoupling was not applied on the $^{1}H$ channel. But the peak at -1.67 ppm showed Gaussian nature in both cases of decoupling and non-decoupling case. Therefore, we can say that the peak at 0.99 ppm has the effect of $^{1}H$-$^{31}P$ heteronuclear coupling, others were not affected by the protons. The line shape fitting and the parameters are shown in Table S1 and Figure S2.

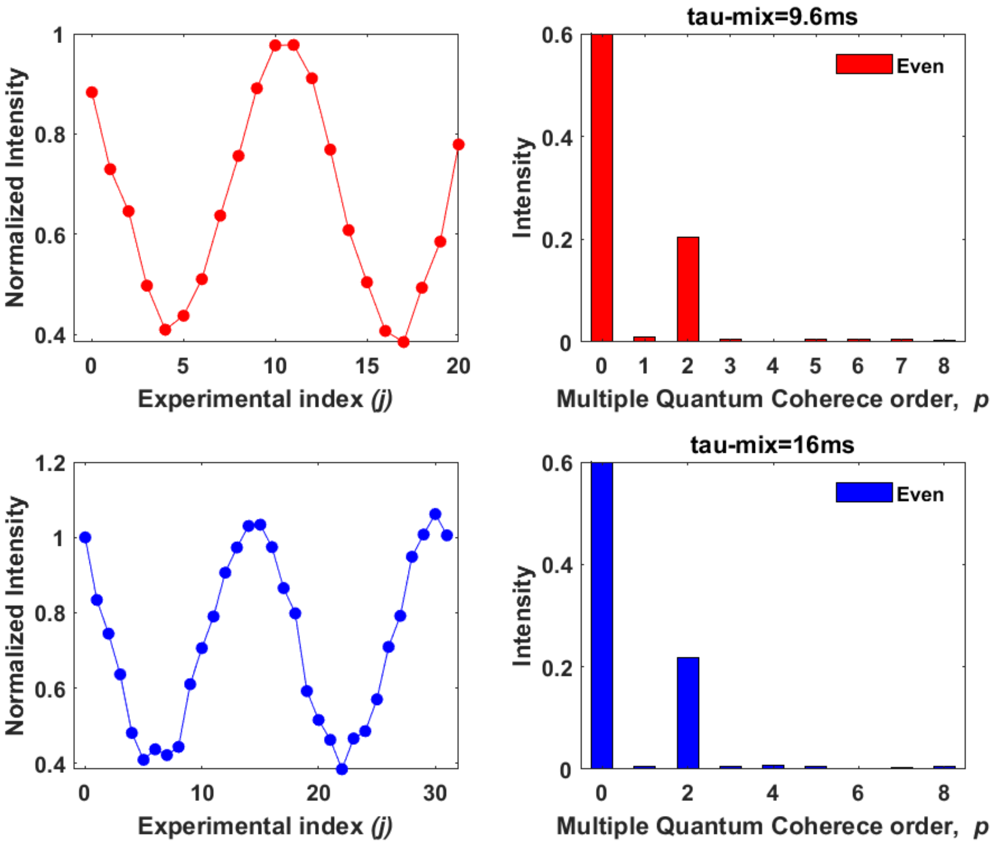

Figure S3. Spin counting experiments with different $SR2^1_8$ mixing times of a) 9.6 ms and b) 16 ms on $^{31}P$ spectra of 1M NaCl S305$^p$ jR2R3 P301L fibrils.

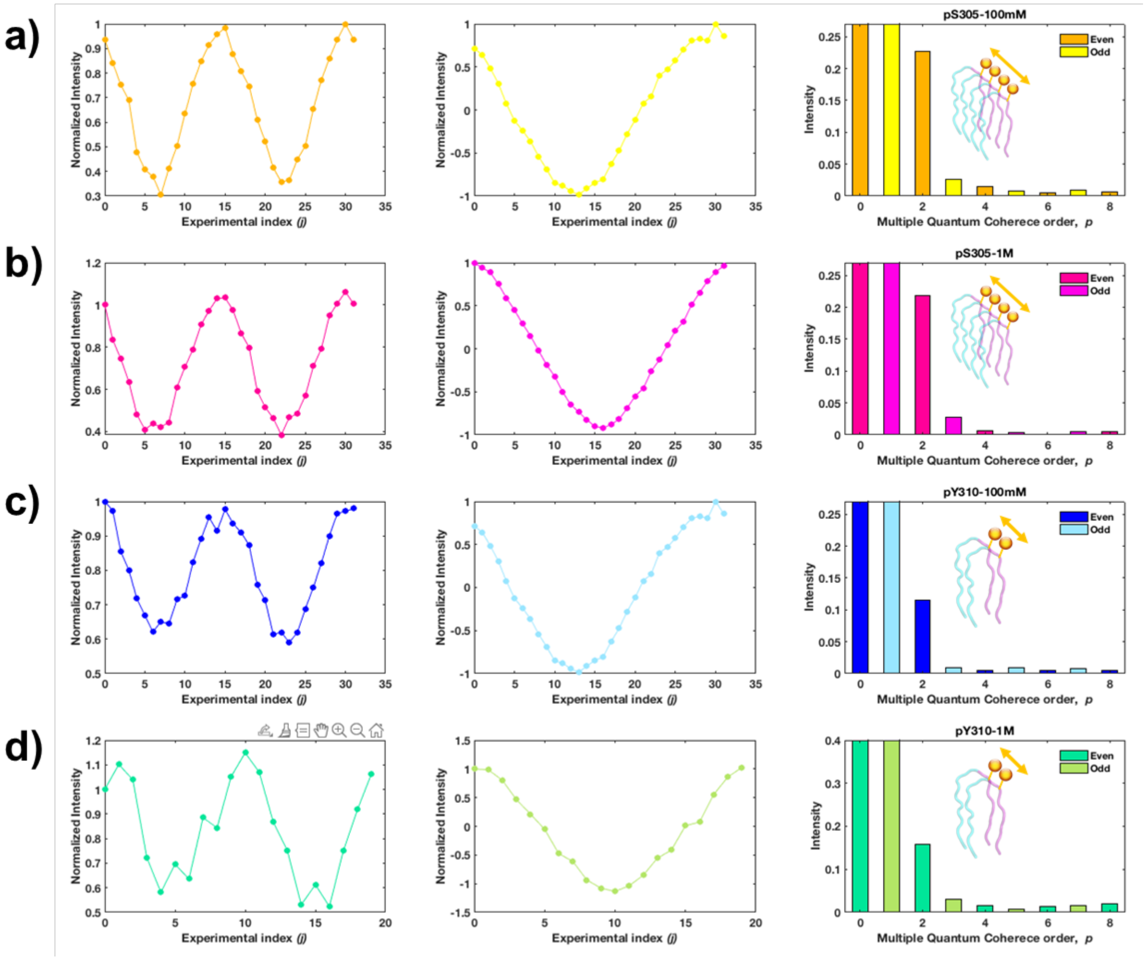

Figure S4. Even and odd spin counting profiles and subsequent Fourier transformed MQCO profiles of S305$^p$ jR2R3 fibrils and Y310$^p$ jR2R3 fibrils at 10 kHz MAS frequency and with 8 ms excitation time during SR2$^1_8$ DQ recoupling sequence under DNP. (a) 100 mM NaCl S305$^p$, (b) 1 M NaCl S305$^p$, (c) 100 mM NaCl Y310$^p$, and (d) 1 M NaCl Y310$^p$.

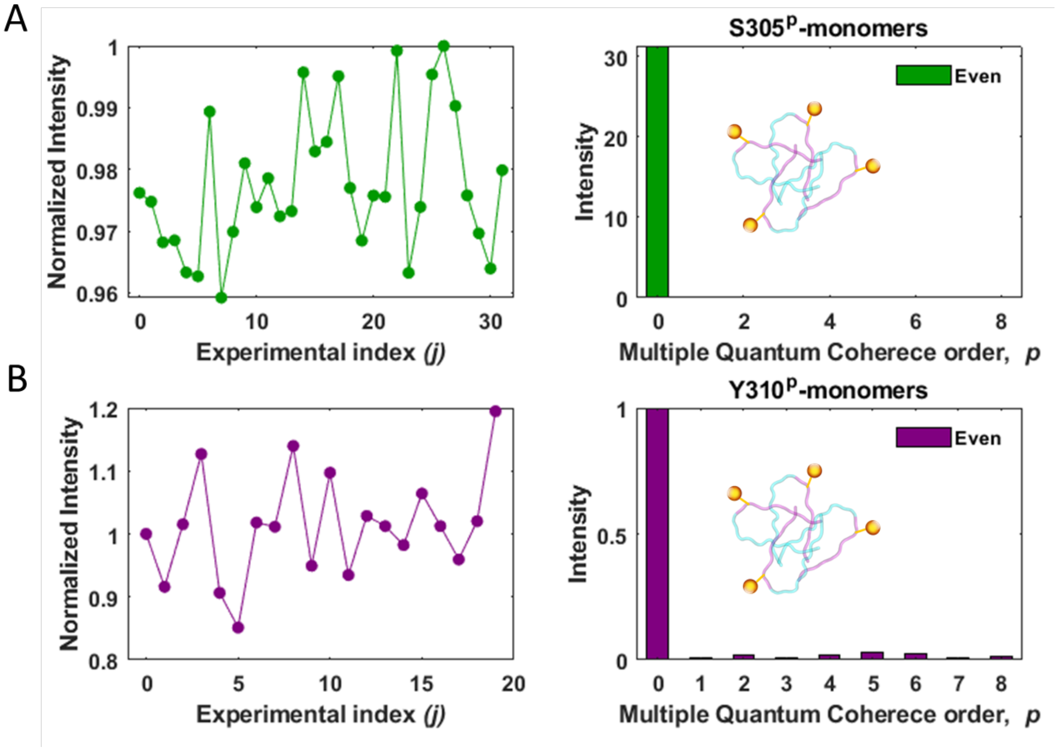

Figure S5. Even MQ-SC profiles and MQCOs extracted from conventional FT method for monomers of S305ᵖ and Y310ᵖ jR2R3 P301L at 10 kHz MAS rate and 100 K temperature under DNP. (A) S305ᵖ jR2R3 P301L monomers and (B) Y310ᵖ jR2R3 P301L monomers. The x-axis of the spin counting profiles is represented by the experimental index (j), where each phase is incremented by 360°/experimental index. All spin-counting profiles' integrals were normalized to the integral of the first experiment (j = 0).

Table S3. Description of the spin systems used for SIMPSON simulations in Figure 6.

| Dipolar | $b_{ij}/(2\pi)$ | $\alpha_{PC}$ | $\beta_{PC}$ | $\gamma_{PC}$ |
|---|---|---|---|---|
| Dimer | | | | |
| dipolar 1 2 | -178.2 | -180.0 | 90.0 | 0.0 |
| dipolar 3 4 | -178.2 | -180.0 | 90.0 | 0.0 |
| dipolar 5 6 | -178.2 | -180.0 | 90.0 | 0.0 |
| Trimer | | | | |
| dipolar 1 2 | -178.2 | -180.0 | 90.0 | 0.0 |
| dipolar 1 3 | -22.3 | -180.0 | 90.0 | 0.0 |
| dipolar 2 3 | -178.2 | -180.0 | 90.0 | 0.0 |
| dipolar 4 5 | -178.2 | -180.0 | 90.0 | 0.0 |
| dipolar 4 6 | -22.3 | -180.0 | 90.0 | 0.0 |
| dipolar 5 6 | -178.2 | -180.0 | 90.0 | 0.0 |
| Six-spin linear | | | | |
| dipolar 1 2 | -178.2 | 179.0 | 97.7 | 0.0 |
| dipolar 1 3 | -22.3 | 179.0 | 97.7 | 0.0 |
| dipolar 1 4 | -6.4 | 179.0 | 97.7 | 0.0 |
| dipolar 1 5 | -3.2 | 179.0 | 97.7 | 0.0 |
| dipolar 1 6 | -1.6 | 179.0 | 97.7 | 0.0 |
| dipolar 2 3 | -178.2 | 179.0 | 97.7 | 0.0 |
| dipolar 2 4 | -22.3 | 179.0 | 97.7 | 0.0 |
| dipolar 2 5 | -6.4 | 179.0 | 97.7 | 0.0 |
| dipolar 2 6 | -3.2 | 179.0 | 97.7 | 0.0 |
| dipolar 3 4 | -178.2 | 179.0 | 97.7 | 0.0 |
| dipolar 3 5 | -22.3 | 179.0 | 97.7 | 0.0 |
| dipolar 3 6 | -6.4 | 179.0 | 97.7 | 0.0 |
| dipolar 4 5 | -178.2 | 179.0 | 97.7 | 0.0 |
| dipolar 4 6 | -22.3 | 179.0 | 97.7 | 0.0 |
| dipolar 5 6 | -178.2 | 179.0 | 97.7 | 0.0 |
| Six-spin jR2R3 | | | | |
| dipolar 1 2 | -167.1 | 90.2 | 90.0 | 0.0 |
| dipolar 1 3 | -20.7 | 90.2 | 90.0 | 0.0 |
| dipolar 1 4 | -202.1 | -179.7 | 81.6 | 0.0 |
| dipolar 1 5 | -63.7 | 133.3 | 84.7 | 0.0 |
| dipolar 1 6 | -15.9 | 115.2 | 86.8 | 0.0 |
| dipolar 2 3 | -167.1 | 90.2 | 90.0 | 0.0 |
| dipolar 2 4 | -65.2 | -132.6 | 84.3 | 0.0 |
| dipolar 2 5 | -198.9 | -179.7 | 82.3 | 0.0 |
| dipolar 2 6 | -65.2 | 133.2 | 84.9 | 0.0 |
| dipolar 3 4 | -15.9 | -114.7 | 86.5 | 0.0 |
| dipolar 3 5 | -63.7 | -132.9 | 84.8 | 0.0 |
| dipolar 3 6 | -202.1 | -179.7 | 82.6 | 0.0 |
| dipolar 4 5 | -167.1 | 90.6 | 90.5 | 0.0 |
| dipolar 4 6 | -20.7 | 90.3 | 90.4 | 0.0 |
| dipolar 5 6 | -167.1 | 89.9 | 90.2 | 0.0 |

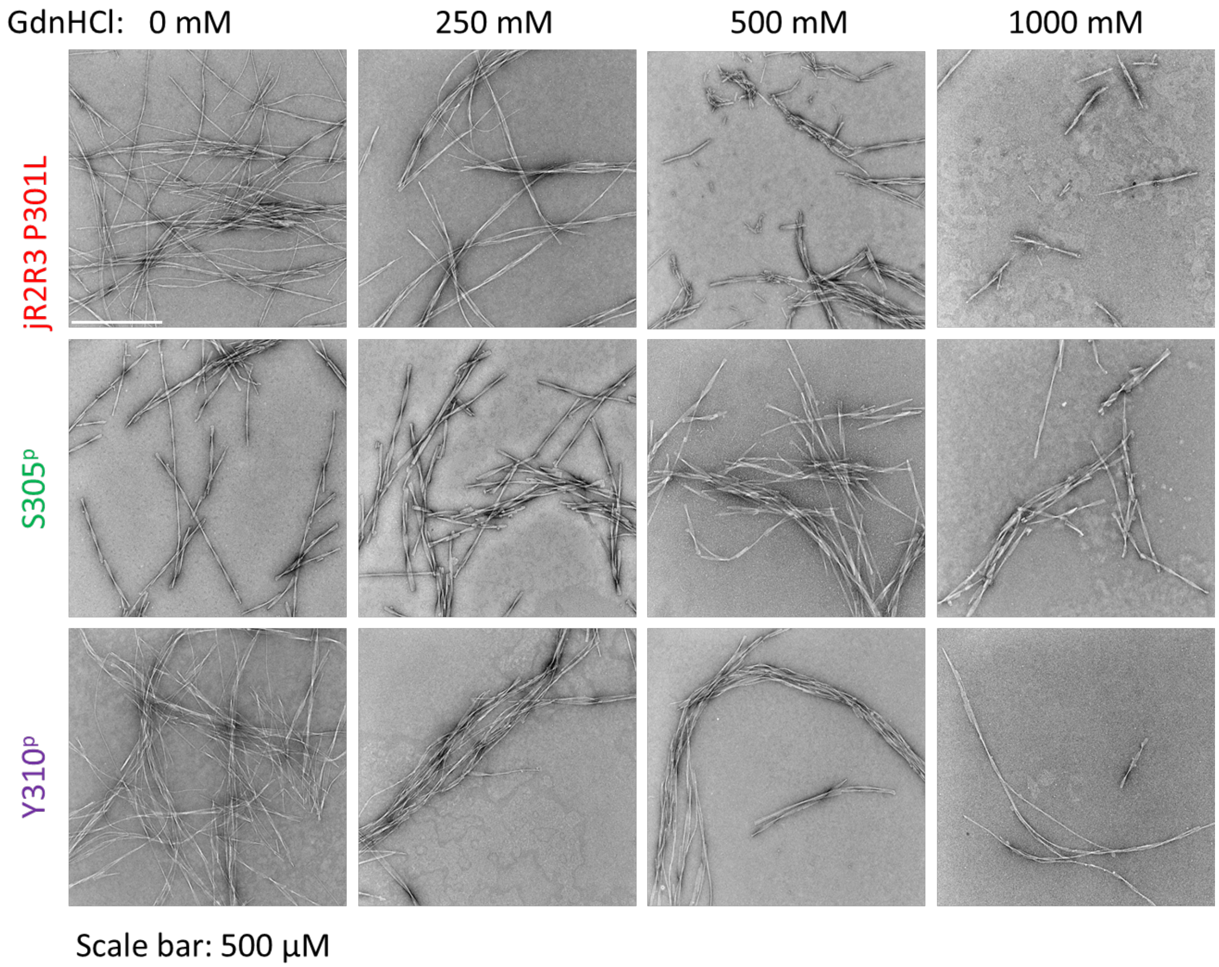

Figure S6. TEM images for stability studies on jR2R3 P301L, S305$^{p}$ and Y310$^{p}$ fibril samples.
